# The orexin-1 receptor antagonist SB-334867 reduces motivation, but not inhibitory control, in a rat stop signal task

**DOI:** 10.1101/467308

**Authors:** Joost Wiskerke, Morgan H. James, Gary Aston-Jones

## Abstract

There is considerable clinical interest in the neuropeptide orexin/hypocretin for its ability to regulate motivation and reward as well as arousal and wakefulness. For instance, antagonists for the orexin-1 receptor (OxR1) are thought to hold great promise for treating drug addiction and disorders associated with overeating, as these compounds repeatedly have been found to suppress seeking of various drugs of abuse as well as highly palatable foods in preclinical models. Given the hypothesized role of OxR1 signaling in cue-driven motivation, an outstanding question is whether pharmacologically blocking this receptor affects cognitive functioning. Response inhibition – the ability to cancel ongoing behavior – is one aspect of cognitive control that may be particularly relevant. Response inhibition deficits are commonly associated with a range of psychiatric disorders and neurological diseases, including substance use disorders and obesity. Moreover, OxR1 signaling recently has been implicated in waiting impulsivity, another aspect of inhibitory control. Here, we investigated the effects of the OxR1 antagonist SB-334867 on response inhibition in a rat version of the stop signal reaction time task. Results show that acutely blocking OxR1 had minimal effects on response inhibition or attentional functioning. In contrast, this manipulation reduced motivation to perform the task and earn food rewards. These results add to the growing body of literature implicating OxR1 in the regulation of motivation and suggest that effects of pharmacological compounds such as SB-334867 on drug seeking behavior are not related to effects on response inhibition.

**Highlights:**

- Orexin-1 receptor antagonists hold great promise for treatment of drug addiction
- These compounds are thought to reduce motivation for drug seeking
- Less is known about effects of orexin-1 receptor blockade on cognitive functioning
- We tested the orexin-1 receptor antagonist SB-334867 in a rat stop signal task
- SB-334867 reduced task motivation but had little effect on executive control as measured with response inhibition

## 1. Introduction

The orexin peptides A and B (hypocretin-1 and 2, respectively) are produced exclusively by neurons of the hypothalamus (Sakurai et al., 1998; deLecea et al., 1998). They exert their actions via two G-protein-coupled receptors (orexin-1 and orexin-2 receptors [OxR1 and OxR2]), which are widely distributed throughout the brain. It is broadly accepted that a dichotomy of function exists between OxR1 and OxR2: signaling at OxR1 is thought to be important for cue-driven reward and motivation, while actions at OxR2 mediate arousal and wakefulness (Brodnik et al., 2018; Mahler et al., 2014). Accordingly, selective OxR1 antagonists reduce seeking of both natural rewards and drugs of abuse, particularly when motivation is augmented by reward-associated stimuli or contexts (James et al., 2017; Mahler et al., 2014). These findings have spurred significant speculation that compounds that interfere with orexin signaling at OxR1 may hold promise as novel treatments to reduce craving and relapse in patients suffering from motivational disorders, including substance use disorders and disorders associated with overeating (Campbell et al., 2018; James et al., 2017; Piccoli et al., 2012; Yeoh et al., 2014).

When considering the potential clinical use of orexin receptor antagonists, it is important to study potential effects of these compounds – both positive and negative – on other processes, such as executive functioning. A crucial part of executive function is response inhibition, which refers to the ability to cancel planned or already initiated actions. Response inhibition is one of the functions required to control impulsivity, or the tendency to engage in behaviors without forethought (Dalley and Robbins, 2017; Evenden, 1999). Impulse control is a complex behavioral construct, which in addition to response inhibition involves decision making and the ability to refrain from acting until it is appropriate (i.e. the ability to wait). The inability to wait, also called “waiting impulsivity”, and deficits in response inhibition are sometimes referred to as forms of motor impulsivity. Much is still unknown about the underlying neurobiology of impulsivity. Importantly however, it is thought that different modalities of impulsivity are mediated by partly distinct neural systems (Broos et al., 2012; Dalley and Robbins, 2017; Eagle and Baunez, 2010; Evenden, 1999; Jentsch et al., 2014).

Recent evidence suggests that OxR1 signaling may mediate certain aspects of impulsivity. The selective OxR1 antagonist SB-334867 (SB) and the dual OxR1/OxR2 antagonist suvorexant have been shown to reduce premature responding in the five choice serial reaction time task (5-CSRTT), a test of waiting impulsivity (Gentile et al., 2017; Muschamp et al., 2014). Moreover, activity of dorsomedial hypothalamus and perifornical area (DMH/PF) orexin neurons was recently found to correlate with successful performance in a Go/NoGo task, a different test of inhibitory motor control (Freeman and Aston-Jones, 2018). In contrast, blockade of orexin signaling with either selective or dual receptor antagonists had no effect in a delay discounting task, which tests impulsive decision making behavior (Gentile et al., 2017). The role of orexin signaling in response inhibition has not been investigated, but it is known that orexin neurons are interconnected with a variety of modulatory neurotransmitter systems thought to be important for response inhibition, including midbrain dopamine neurons, norepinephrine neurons in the locus coeruleus, and basal forebrain cholinergic neurons (Balcita-Pedicino and Sesack, 2007; Espana et al., 2005; Fadel and Deutch, 2002; Fadel and Burk, 2010; Vittoz and Berridge, 2006). Orexin signaling has also been shown to modulate glutamatergic thalamocortical synapses that ultimately alter acetylcholine and glutamate release in the prefrontal cortex (Calva and Fadel, 2018; Lambe et al., 2005), a part of the brain thought to be crucial for inhibitory control. Finally, substance use disorder is characterized by high levels of impulsivity, including response inhibition (Argyriou et al., 2018; Jentsch et al., 2014; Verdejo-Garcia et al., 2008), perhaps pointing to a common role for the orexin system in mediating these processes.

Response inhibition is commonly measured in clinical populations using a stop signal task (SST). In this task, subjects perform a rapid motor response to a stimulus (Go signal) to earn a reward. On some trials the Go stimulus is followed by a distinct stimulus (Stop signal) and the subject must inhibit performance of the Go response (Logan, 1994). Following its initial translation to a rat model (Eagle and Robbins, 2003; Feola et al., 2000), SST procedures for rats and mice have in recent years been expanded and refined, with the aim to identify neural circuits underlying response inhibition (Pattij et al., 2009; Bryden et al., 2012; Leventhal et al., 2012; Humby et al., 2013; Maguire et al., 2018; Mayse et al., 2014). In these tasks, rodents are typically food- or water-restricted and are rewarded with a small food/liquid reward for correct “going” and “stopping” behavior. Therefore, in addition to measuring response inhibition and attention, it is possible to derive tentative measures of food motivation from indices such as trial omission rate and latency to collect food rewards from the feeder (e.g. (Bari and Robbins, 2013; Humby et al., 2013). In this study, we administered the selective OxR1 antagonist SB to rats and examined its effects on performance in the SST to investigate the role of OxR1 signaling in response inhibition.

## 2. Results

In this study, we characterized the effects of acute administration of the OxR1 receptor antagonist SB in a rat version of the SST (Figure 1).

**Figure 1.**
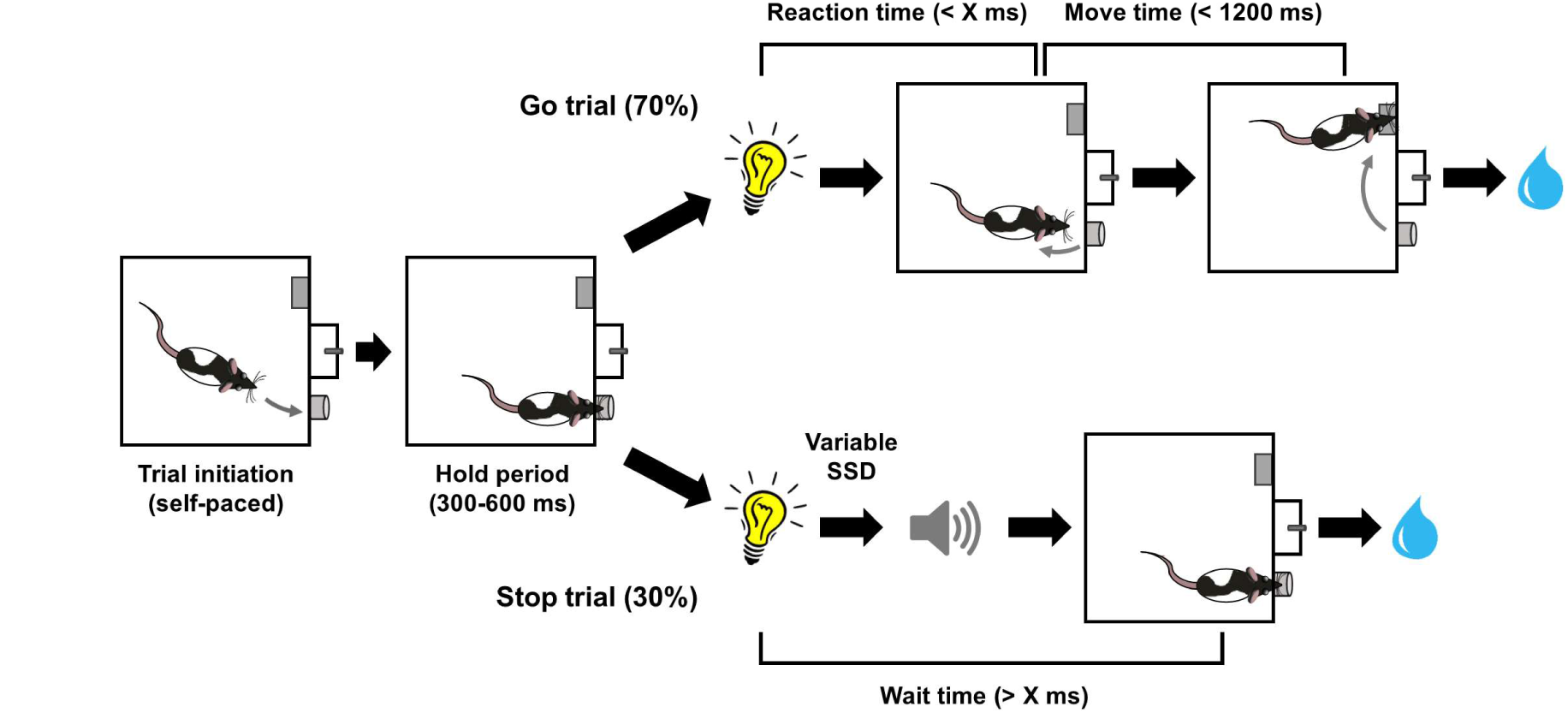
Schematic representation of the rat stop signal task. Flow of events during Go and Stop trials. “X” on Go trials represents the maximum reaction time allowed for exiting the nosepoke port following presentation of the Go cue. The same cut off value was used as the minimum time the rat had to maintain fixation in the nosepoke port after the Go cue presentation on Stop trials. Prior to the first test day, X was individually titrated per rat to a value between 400-700 ms. The stop signal delay (SSD) was variable across Stop trials. While correctly executed Go and Stop trials were rewarded with 60 μl of 15% sucrose water, incorrect performance on Go (Reaction time > X ms or Move time > 1200 ms) and Stop trials (Wait time < X ms) resulted in a 5 s time out during which the house light was turned on.

### 2.1 Effects of a vehicle injection

We compared the behavioral effects of two doses of SB (10 and 30 mg/kg) relative to an injection with its vehicle (2% dimethyl sulfoxide, 10% 2-hydropropyl-β-cyclodextrin in sterile water). We first checked for putative behavioral effects of the vehicle itself by comparing its effects with a measure of baseline behavioral performance (average performance over three training sessions), looking at four main behavioral parameters of the SST. Pearson correlational analyses showed that vehicle and baseline results correlated well for probability of a correct Go trial (*p*(Correct Go); R^2^ = 0.53, adjusted *p* < 0.001) and median Go trial reaction time (GoRT; R^2^ = 0.76, adjusted *p* < 0.001). In contrast, weaker correlations were found for two measures of stop performance: estimated stop signal reaction time (SSRT; R^2^ = 0.29, adjusted *p* = 0.05) and *p*(Failed Stop), the probability of failing to inhibit the Go response on Stop trials (R^2^ = 0.23, adjusted *p* = 0.11). Thus, we decided to include baseline performance as an additional control condition in all subsequent analyses.

### 2.2 SB-334867 did not affect responding during Go and Stop trials

We examined the effects of acute administration of SB on measures of performance on Go and Stop trials to test whether blocking OxR1 may affect response inhibition as measured in the SST (Figure 2). Because we planned to look at many behavioral parameters, we decided to use a stringent significance threshold (*p* < 0.005) for the repeated measures analyses of variance, while Bonferroni correction was used for any *post hoc* comparisons (with *p* < 0.05 as significance threshold).

**Figure 2.**
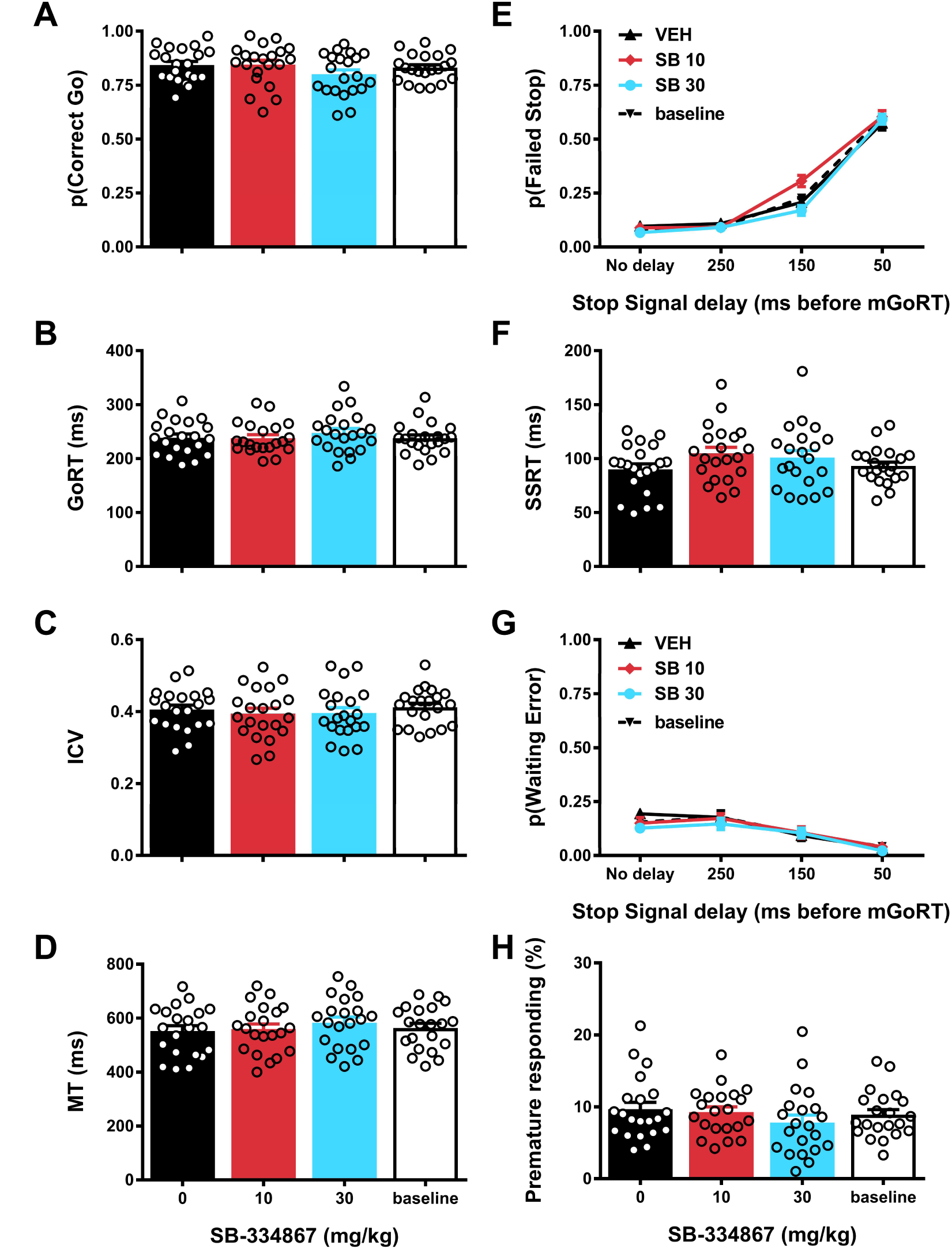
Effects of SB-334867 on performance during Go and Stop trials in the stop signal task. Effects of acute administration of the OxR1 receptor antagonist SB-334867 or its vehicle on probability of correct Go trials (A), median reaction time on correct Go trials (B), the ICV or intra-individual coefficient of variation [standard deviation GoRT/mean GoRT] (C), median time between exit of the nosepoke port and a lever press on correct Go trials (MT, D), probability of failed Stop trials (E), estimated stop signal reaction time (SSRT; F), probability of waiting errors during Stop trials (G) and percentage of trials with a nosepoke port exit during the hold period (H). For the baseline measurement, data from the training sessions immediately preceding the 3 test days was averaged. Error bars represent standard error of the mean, while open circles represent data from individual rats (N=21).

Performance on Go trials was analyzed first, to check for drug effects on attentional functioning, reaction speed and motor activity. Effects on Go trial performance would indicate disruption of general task performance, which would preclude investigating SB effects on response inhibition. Although the results revealed a trend towards a treatment effect on the Go trial accuracy measure *p*(Correct Go), this did not reach the significance threshold of *p* < 0.005 (Figure 2a; F_3,60_ = 3.24, *p* = 0.03). Similarly, no significant effects were observed for GoRT, the intra-individual coefficient of variation (ICV, a measure of reaction time variability; (e.g.(Bellgrove et al., 2004)) or movement time (MT, measure of movement speed/locomotor activity) (Figure 2b-d; median GoRT: F_3,60_ = 2.23, *p* = 0.09; ICV: F_3,60_ = 1.17, *p* = 0.33; MT: F_3,60_ = 3.12, *p* = 0.05, ε = 0.74).

After we had established that SB administration did not disrupt ability to perform Go trials, we switched our attention to Stop trial performance to assess putative effects of SB on inhibitory control. Successful performance of Stop trials in the SST requires two partly distinct aspects of inhibitory control: response inhibition (also called action cancelation) and the ability to wait (also called action restraint). We used the recently-developed “modified integration method” to disentangle these two subdomains of inhibitory control (Mayse et al., 2014). With this method, response inhibition can be assessed by calculating *p*(Failed Stop) and SSRT. Our results showed that SB administration affected neither measure, although the interaction effect between drug treatment and stop signal delay (SSD) for *p*(Failed Stop) almost reached statistical significance (Figure 2e-f; Treatment effect *p*(Failed Stop): F_3,60_ = 3.66, *p* = 0.02; Treatment × SSD effect *p*(Failed Stop): F_9,180_ = 3.63, *p* = 0.007, ε = 0.54; Treatment effect SSRT: F_3,60_ = 2.20, *p* = 0.10). To test for SB effects on waiting impulsivity, we first examined the parameter *p*(Waiting Error). This refers to the proportion of Stop trials on which a rat initially succeeds to cancel the Go response but subsequently fails to wait until the offset of the Stop tone. The data showed no significant SB effects on this measure (Figure 2g): Treatment effect: F_3,60_ = 1.99, *p* = 0.12; Treatment × SSD effect: F_9,180_ = 1.01, *p* = 0.43. Similarly, we found no significant effects of SB on a second measure of waiting impulsivity, premature responding, i.e. the percentage of trials (Go and Stop trials combined) that rats prematurely exited the nosepoke port during the variable hold period prior to the onset of the Go cue (Figure 2h; F_3,60_ = 3.77, *p* = 0.03, ε = 0.64). Thus, we found no evidence of robust effects of SB on either response inhibition or the ability to wait as required to perform the SST.

To further rule out that SB affected Go and/or Stop trial performance, we additionally obtained signal-detection-based measures of trial type discriminability and overall response bias (Macmillan and Creelman, 2005; Stanislaw and Todorov, 1999; Swets et al., 1961). In the context of our study, the measure d’ represents a subject’s ability to discriminate between the two stimulus-response options in the SST. Thus, this value would be highest if a subject executed the Go response (exit nosepoke port and press the lever) only on Go trials, and the Stop response (stay in the nosepoke port) exclusively on Stop trials. The measure c (bias) represents the tendency of a subject to execute either the Go response or the Stop response, independent of trial type. Results of these analyses showed that SB did not affect the value obtained for d’ (Figure 3a; F_3,60_ = 1.47, *p* = 0.23). On the other hand, there was a significant treatment effect on the value obtained for c (Figure 3b; F_3,60_ = 6.78, *p* < 0.001). *Post hoc* testing indicated that compared to vehicle, administration of 30 mg/kg SB changed response bias towards “staying” (i.e. lower c-value). It should be noted that this drug effect might be related to the fact that rats showed a significantly larger bias towards “going” under vehicle conditions as compared to baseline. Together, the data indicate that SB administration had minimal, if any, effects on Go and Stop trial performance in our SST.

**Figure 3.**
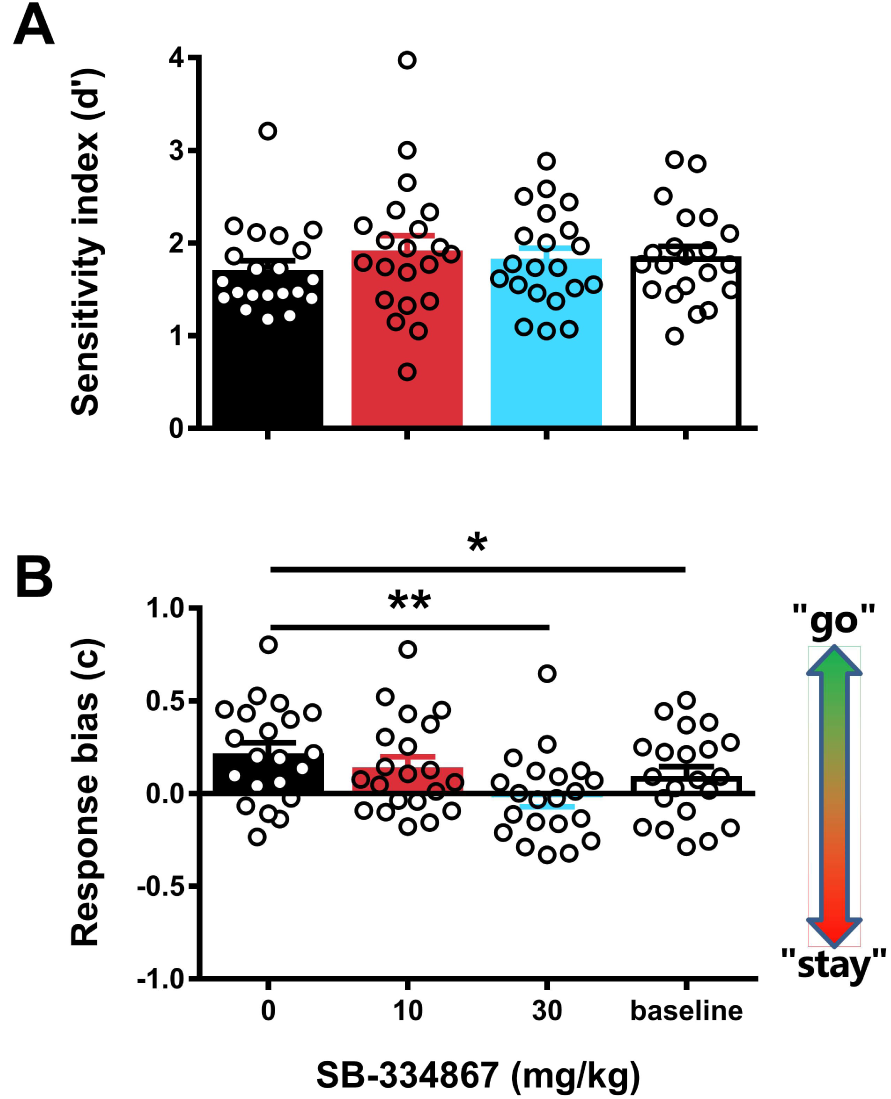
Effects of SB-334867 on cue discrimination and response bias in the stop signal task. Effects of acute administration of the OxR1 antagonist SB-334867 or its vehicle on signal detection theory measures of cue discrimination (A) and response bias across all trials (B). For the baseline measurement, data from the training sessions immediately preceding the 3 test days was averaged. Error bars represent standard error of the mean, while open circles represent data from individual rats (N=21). ^*^ p < 0.05, ^**^ p < 0.005 as compared to vehicle treatment.

### 2.3 SB-334867 reduced motivation and task engagement

A change in response bias towards “staying” could be an indication of reduced motivation to perform the task. A lack of motivation is likely to result in a tendency to more frequently opt for the less effortful “stay” option, irrespective of the stimuli that are presented during the trial. To further investigate potential motivation-reducing effects of SB in the SST, we first examined the rats’ pace in the task as reflected by the average number of trials executed per minute (Figure 4a). Results showed a significant treatment effect (F_3,60_ = 14.45, *p* < 0.001, ε = 0.69) with further analysis showing that administration of 30 mg/kg SB reduced the trial execution rate. Next, we determined whether this effect on trial execution rate was caused by rats taking long breaks during the task, or whether it was due to a more subtle but consistent lengthening of the inter-trial interval. We found a significant treatment effect on omission rate (Figure 4b; F_3,60_ = 13.61, *p* < 0.001), caused by an increase in trial omissions following administration of the higher dose of SB. Finally, we analyzed the data on feeder latencies, i.e. the time to collect a reward after successful trial completion. This parameter is thought to reflect a measure of motivation to obtain reward (e.g. (Bari and Robbins, 2013; Humby et al., 2013). Results showed a significant overall treatment effect (Figure 4c; F_3,60_ = 8.75, *p* < 0.001), and again *post hoc* testing revealed that 30 mg/kg SB resulted in an increased median latency to collect rewards, relative to vehicle treatment. Together with the other results, these findings are consistent with SB reducing motivation to perform in the SST while minimally affecting response inhibition and attentional functioning.

**Figure 4.**
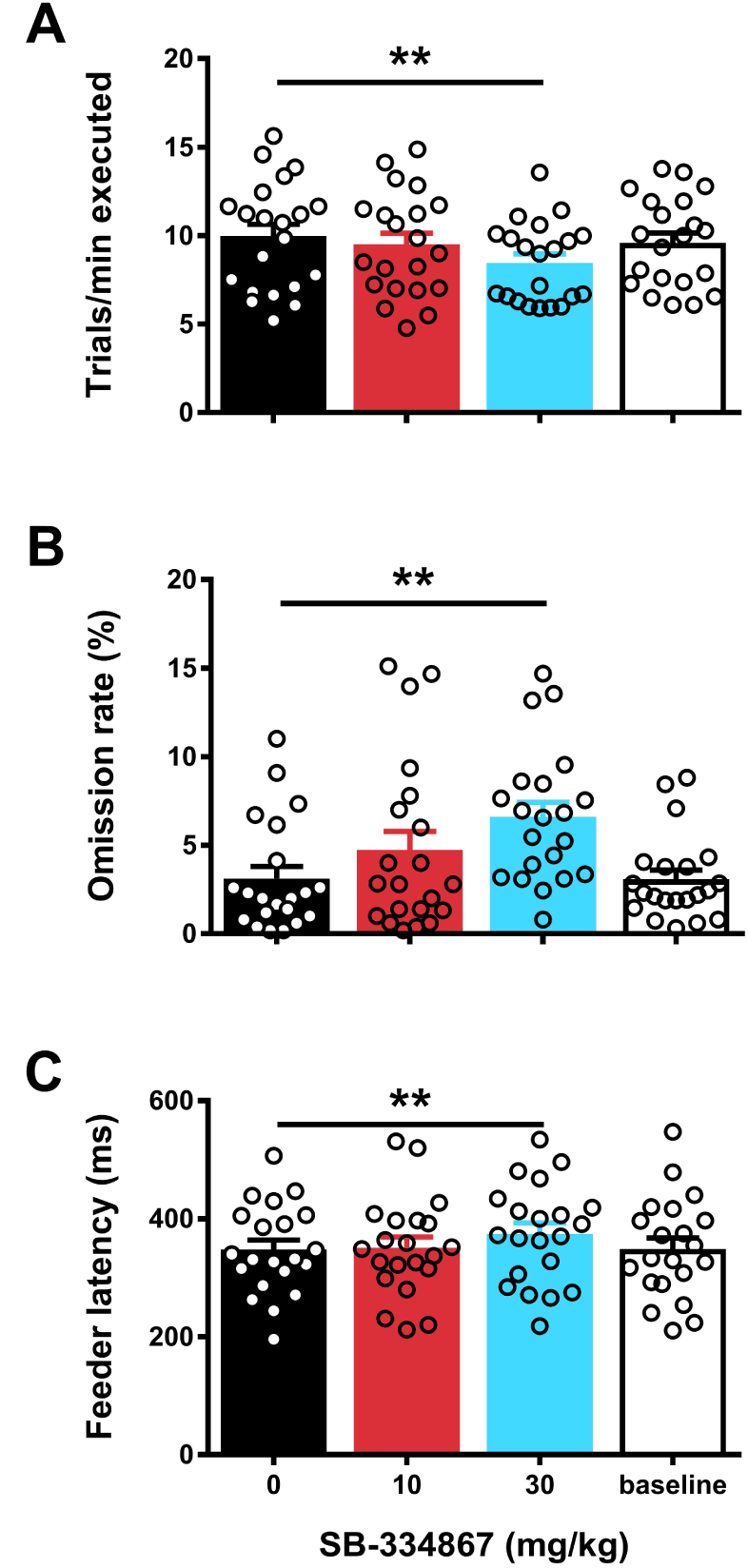
Effects of SB-334867 on motivational parameters in the stop signal task. Effects of acute administration of the OxR1 antagonist SB-334867 or its vehicle on the rate of trial execution (A), the *%* of trials omitted (B) and the mean latency to collect rewards during correct trials (C). For the baseline measurement, data from the training sessions immediately preceding the 3 test days was averaged. Error bars represent standard error of the mean, while open circles represent data from individual rats (N=21). ^**^ Reflect p < 0.005 as compared to vehicle treatment.

## 3. Discussion

The hypothalamic orexin neuropeptide system has been shown to be a critical regulator of cue-driven motivation for natural and drug rewards (for review see e.g. (James et al., 2017; Mahler et al., 2014)). Less is known about its role in top-down cognitive processes such as impulse control. Here, we examined the role of the orexin system in mediating performance in a rat version of the SST, a task that requires rats to exhibit behavioral control in response to discrete cues to earn sucrose rewards. We show that pretreatment with the selective OxR1 antagonist SB had no significant effects on measures of response inhibition or the ability to wait in the SST. However, we observed SB-induced reductions in several measures that reflected motivation of the rats to perform the task and earn food rewards. Thus, our data indicate that, despite a clear role for OxR1 signaling in cue-driven reward seeking and motivation, this neuropeptide system does not appear to be critical for stimulus-driven inhibitory control.

### 3.1 Orexin signaling and inhibitory control

We used the SST to assess two distinct aspects of inhibitory control: response inhibition and waiting impulsivity. Response inhibition was assessed by examining ability of the rats to correctly inhibit an already initiated action in response to a Stop signal across several SSDs, as well as by estimating rats’ SSRTs. Pretreatment with SB at 10 or 30mg/kg had no significant effect on either measure. SB also did not affect waiting behavior, measured by the ability to appropriately delay exiting the nosepoke port. Specifically, SB administration did not affect the rate of premature exits from the nosepoke before presentation of the Go stimulus, nor did it affect the percentage of waiting errors made during stop trials.

These findings are surprising for a few reasons. First, Ox1R signaling has been implicated in several psychiatric disorders that are commonly associated with deficient impulse control. Most notably, addiction is associated with higher numbers, activity and excitability of orexin cells (James et al., 2018b; Lawrence et al., 2006; Moorman et al., 2016; Thannickal et al., 2018; Yeoh et al., 2012; Yeoh et al., 2018), and SB is highly effective at reducing several addiction-related behaviors in rodents (Brodnik et al., 2018; Campbell et al., 2018; James et al., 2017; James et al., 2018a; Martin-Fardon and Weiss, 2014a; Martin-Fardon and Weiss, 2014b; Moorman et al., 2017; Perrey and Zhang, 2018; Schmeichel et al., 2017). Secondly, orexin neurons are interconnected with a variety of brain regions (e.g. prefrontal cortex) and modulatory neurotransmitter systems (e.g. dopamine, norepinephrine and basal forebrain acetylcholine) thought to be important for arousal, attention and impulsivity (Balcita-Pedicino and Sesack, 2007; Calva and Fadel, 2018; Espana et al., 2005; Fadel and Deutch, 2002; Fadel and Burk, 2010; Lambe et al., 2005; Vittoz and Berridge, 2006). Thirdly, and perhaps most relevant in this case, several recent studies have implicated orexin signaling in other forms of motor impulsivity. For example, our lab recently reported that activation of the DMH/PF, but not LH, orexin neurons was positively correlated with successful performance in a Go/NoGo task (Freeman and Aston-Jones, 2018), a task in which rats must inhibit a prepotent Go response on a subset of trials in which a No-Go stimulus is presented instead of the Go stimulus. More direct evidence supporting a role for orexin signaling in regulating inhibitory control has come from two studies using a 5-CSRTT, an operant task in which subjects are required to withhold responding until a stimulus light is illuminated in one of five locations. Acute administration of SB or the dual orexin receptor antagonist suvorexant in this task reduced the number of premature responses without affecting accuracy, omission rate or feeder latency (Gentile et al., 2017; Muschamp et al., 2014). These results are in contrast with our findings that SB did not significantly affect response inhibition or premature responding but did increase omission rate and feeder latencies. Importantly, 5-CSRTT and Go/NoGo tasks measure aspects of motor impulsivity that are distinct from the response inhibition measured in a SST as used here (for review, see e.g. (Eagle and Baunez, 2010). The discrepancies between the effects of SB on premature responding, omission rate and feeder latency in the 5-CSRTT and SST could also be related to differences in task design. The SST puts a premium on very fast responding, consists of hundreds of trials per session, and requires subjects to withhold responding only for subsecond periods at a time. The 5-CSRTT, on the other hand, places much higher demands on spatial attention, consists only of around a hundred trials per session, and requires action restraint during 5 s inter-trial intervals. It should also be noted that the studies by Muschamp and co-workers used a fixed duration for the pre-stimulus waiting period in the 5-CSRTT (Gentile et al., 2017; Muschamp et al., 2014). Consequently, it cannot be ruled out that the reported effects of SB and suvorexant on premature responding in those studies reflected an improvement in time estimation rather than inhibitory control. Altogether, the data attest to the multifaceted nature of behavioral constructs such as inhibitory control and the importance that even subtle differences in task design may have on results.

Another conclusion from our data may be that if orexin signaling modulates inhibitory control, it does so only under specific conditions. Here we did not assess the effects of SB on drug-induced impulsivity or response inhibition in subjects with a history of drug or palatable food intake. It may be interesting to investigate this given that OxR1 antagonists have been shown to block cocaine-induced increases in impulsivity in the 5-CSRTT (Gentile et al., 2017; Muschamp et al., 2014). Recent evidence from our lab and others also demonstrated that drug exposure or a history of high fat food intake can increase the number and excitability of orexin-expressing neurons (James et al., 2018b; Lawrence et al., 2006; Park et al., 2004; Thannickal et al., 2018; Yeoh et al., 2012; Yeoh et al., 2018). Thus, it cannot be ruled out that the role of orexin signaling in executive functions such as response inhibition changes following exposure to drugs of abuse or following a period of overconsumption of palatable food.

### 3.2 Orexin signaling and motivation

We observed several behavioral consequences of SB pretreatment on SST task performance that most likely reflected attenuated motivation to seek sucrose reward. First, following SB treatment, rats exhibited a stronger bias towards “staying” (i.e. not responding to the Go cue) regardless of trial type. Second, rats engaged in fewer trials per minute and registered more trial omissions following pretreatment with SB. Although increased omissions could be interpreted as an attentional deficit, this is inconsistent with the fact that SB had no effect on Go or Stop trial accuracies, trial type discrimination (i.e., d’-value) or ICV (measure of attentional lapses, e.g. (Bellgrove et al., 2004)). Third, on trials where rats were rewarded with a sucrose reward following correct Go or Stop behavior, rats exhibited a greater latency to collect this reward following 30 mg/kg SB pretreatment. Importantly, we do not believe that these motivational effects are attributable to sedative effects of SB treatment, for two reasons. First, on successful Go trials, SB had no effect on time taken to leave the nosepoke port following a Go signal (i.e. GoRT) or time to press the lever following exiting the nosepoke port upon presentation of the Go signal (i.e. MT). These two findings indicate that the rats’ ability to execute the task quickly was not impaired following SB administration. Second, we and others have previously shown that SB at doses used here has little or no effect on motor behavior as assessed by measurement of total distance traveled in a test of general locomotor activity (James et al., 2018b; Porter-Stransky et al., 2015; Smith et al., 2009b) or low-effort responding for food and drug rewards (Bentzley and Aston-Jones, 2015; Boutrel et al., 2005; Espana et al., 2010; Hutcheson et al., 2011; James et al., 2018b; Smith et al., 2009b). Thus, the most parsimonious explanation of the behavioral effects of SB in the current study is a reduction in motivation of the rats to work for a palatable food.

In the present study, rats were food-restricted to 85-90% of free-fed intake and thus, under baseline conditions, they were highly motivated to successfully execute trials to earn liquid sucrose rewards. This is evidenced by the high number of trials completed (averaging ~10 trials/minute across a session) and the large amount of sucrose water earned throughout the session (routinely 15-20 mL/session). Our findings therefore are in line with a large body of evidence pointing to a role for orexin signaling in the seeking of drugs or food associated with high motivational valence, rather than their simple consumption, *per se* (Freeman and Aston-Jones, 2018; James et al., 2017). For example, SB has no effect on cocaine self-administration on a low-effort schedule of reinforcement (fixed ratio 1), but reliably decreases high-effort (fixed ratio 5 or higher) and progressive ratio responding for this reinforcer (Borgland et al., 2009; Hollander et al., 2012; Smith et al., 2009a). Similarly, SB reduces effortful seeking as well as binge-like intake of foods rich in fat and/or sugar – and non-caloric palatable substances such as saccharin – at doses that do not affect regular chow intake (Alcaraz-Iborra et al., 2014; Borgland et al., 2009; Choi et al., 2010; Nair et al., 2008; Olney et al., 2015; Rorabaugh et al., 2014; Valdivia et al., 2014; Valdivia et al., 2015; Vickers et al., 2015). The orexin system appears to be particularly important for driving reward seeking under circumstances where motivation is enhanced by the presence of external stimuli. Both systemically and intracranially-administered SB reliably blocks reinstatement of both drug and food seeking elicited by discrete stimuli and/or contexts previously associated with reward availability (Cason and Aston-Jones, 2013a; Cason and Aston-Jones, 2013b; James et al., 2011; James et al., 2018b; Martin-Fardon and Weiss, 2014a; Moorman et al., 2017; Smith et al., 2009a; Smith et al., 2010). Our current data indicate that these effects of SB on cue-driven reward seeking are due to a specific role for the orexin system in modulating the motivation-imbuing properties of the reward stimuli rather than response inhibition or attentional functioning, although it will be important for future studies to confirm this. Such studies should keep in mind that, despite being the most widely used antagonist to test Ox1R function, SB’s selectivity for OxR1 over OxR2 is modest (~50-fold selectivity) (Porter et al., 2001; Smart et al., 2001). SB also has some affinity for other G-protein coupled receptors, including serotonin 2B and 2C receptors (Porter et al., 2001; Smart et al., 2001). Several alternative compounds with greater selectivity for OxR1 have recently been developed (Perrey and Zhang, 2018) and could be tested in the SST and other assays of impulsivity to further characterize the role for OxR1 signaling in this aspect of cognitive behavior.

### 3.3 Concluding remarks

We showed that the OxR1 antagonist SB has minimal effects on response inhibition or attentional functioning in a rat version of the SST. In contrast, SB reduced motivation to perform the task and earn sucrose rewards. These findings add to a growing body of literature implicating signaling at OxR1 in the regulation of cue-driven motivation for reward. Confirmation of a selective role for the orexin system in mediating motivational processes is particularly timely given recent discussion surrounding the potential utility of orexin-based compounds for the treatment of motivational disorders, including drug addiction and binge-eating disorder. Thus, while OxR1 antagonists appear useful for reducing stimulus-driven craving in patients with such psychiatric disorders, the utility of these compounds in decreasing symptoms of comorbid impulsivity might be limited. At the same time, our SST data, together with previous data in the 5-CSRTT and delay discounting task (Gentile et al., 2017; Muschamp et al., 2014), suggest that OxR1 antagonist treatment is unlikely to overtly affect cognitive functioning as a side effect. To further validate the clinical potential of OxR1 antagonists, the effects of chronic administration of such compounds on cognitive performance should be studied in more detail.

## 4. Experimental Procedures

### 4.1 Animals and husbandry

Twenty-seven male Long Evans rats (Charles River, Kingston, NY), weighing 300–400 g at the start of the experiment, were used, divided into two cohorts. Upon arrival, animals were pair-housed in a temperature- and humidity-controlled animal facility (AAALAC-accredited) in clear, plastic IVC caging (Allentown Inc., USA) with *ad libitum* access to water and regular chow (PicoLab Rodent Diet 20 (5053), LabDiet, St. Louis, MO). Throughout the entire experiment, animals were kept on a reverse 12-hour light cycle (lights off 8.30 am), with experimental sessions taking place 5-7 days/week between 10 am and 3 pm. For motivational purposes, after an acclimation period of 7 days and 1-2 days prior to the first training session, rats were food-restricted to 85-90% of free-fed intake. Rats remained food-restricted until the end of the experiment, with a daily opportunity to supplement calorie intake through sucrose rewards in the operant task. Water was available *ad libitum* in the homecage throughout the entire experiment. Animals remained pair-housed during the initial training stages but were single-housed for a minimum of two weeks before the first test day to gain better control over food intake of individual rats. All procedures were approved by the Rutgers University Institutional Animal Care and Use Committee.

### 4.2 Apparatus

Experiments were conducted in 8 standard rat operant chambers with stainless steel grid floors (Med Associates Inc., St. Albans, VT) housed in sound-insulating and ventilated cubicles. Each chamber was equipped with one retractable lever and one nosepoke port on the same wall, separated by a central reward port containing a liquid receptacle and a yellow cue light in its ceiling. The locations of the lever and nose poke port relative to the reward port were counterbalanced across operant chambers. Customized triple light emitting diode (LED) stimulus displays (ENV-222M) were located directly above the nosepoke port and the lever. Only the centered red LED was used above the nosepoke port, while only the two outside green LEDs were used above the lever. The nosepoke port additionally contained an internal yellow LED light. Both the nosepoke port and the reward port contained infrared detectors to detect entries. A speaker was positioned at the top of the wall above the nosepoke port. Finally, a white house light was centered at the top of the opposite chamber wall. Sucrose water could be delivered to the liquid receptacle via a syringe pump located on top of the operant chamber (inside the sound-attenuating chamber). A Windows computer equipped with MED-PC IV (Med Associates Inc. St. Albans, VT) controlled experimental sessions and recorded data with a 1 ms time resolution.

### 4.3 Behavioral experiment

Shaping and stabilizing of the rats’ behavior in our SST (Figure 1) took about 2 months (~50 training sessions). Although the exact training trajectory was individualized for each rat, it always consisted of the same four phases that are described below.

#### 4.3.1 Phase 1: Habituation

The first step of the training was an overnight program (15 hours or 900 trials per session, whichever came first). During the first 100 trials, rats learned to collect sucrose rewards (60 μl 15% sucrose water) by poking their nose into the reward port. The cue light inside the reward port was briefly turned on during reward delivery. During trials 101-200, rats were required to first make a response in either the nosepoke port or on the lever before they could collect a sucrose reward in the reward port. Opportunities to nosepoke or lever press from now on were cued by the stimulus lights above the nosepoke and the lever, while reward availability was cued by onset of the light inside the reward port. During trials 201-300, rats had to alternate between nosepoking and lever pressing to obtain a reward, to assure that all rats became familiar with both response units. The stimulus lights above the nosepoke and lever indicated which response had to be made. Then rats entered the final stage of habituation (trial 301 and onwards), in which they were trained to maintain the nosepoke for increasing durations before lever pressing to get a reward. The required minimum hold period increased in steps of 50 ms per 10 correct trials, up to a maximum of 1 s. The required minimum hold period was cued by the onset of the stimulus light inside the nosepoke port upon the rat entering it followed by the offset of the same light once the minimum hold period had passed. Premature exits from the nosepoke port resulted in a 5 s timeout during which the house light was turned on. This final stage lasted a minimum of 300 trials. Rats were transferred to the next training phase when they could reliably stay in the nosepoke port for ≥ 1 s before making the lever press (> 80% accuracy for ≥ 100 trials). If a rat did not reach this criterion during the first overnight session, a second habituation session was conducted. That session was then started at the final training stage reached by that rat during the first session.

#### 4.3.2 Phase 2: Learning to execute Go trials fast and correctly

During the second step of the training, rats learned to execute Go trials as quickly as possible. Sessions occurred during the day and session duration was decreased to 120 min. Each trial started with a variable nosepoke hold time (300-600 ms), after which the light inside the nosepoke port turned off and the stimulus lights above the lever turned on. Upon presentation of this Go cue, the rat had to exit the nosepoke port as quickly as possible to go and press the lever on the other side of the wall and get a sucrose reward. Rats started off with 60 s limits for both the time to exit the nosepoke port (i.e GoRT) and the time to press the lever (i.e. MT). These time limits were gradually reduced as rats improved performance. When the GoRT limit had reached 1.5 s and the MT limit 3 s, we added discouragement of random responding. Lever presses made before entering the nosepoke port at the start of a trial, or reward port entries made in between exiting the nosepoke port and pressing the lever now resulted in a 5 s timeout. Simultaneously, we also started registering trial omissions, with failures to initiate the next trial within 60 s after finishing a trial resulting in a timeout period with the house light on. Since we consider such omissions a reflection of task disengagement, rats were required to make an operant response (lever press or nosepoke) to actively end the timeout (indicated by house light being turned off and the red cue light above the nosepoke port turning back on) and get the opportunity to initiate a new trial. Rats transitioned to the next phase in the training when their GoRT limit was 1-1.2 s and their MT limit was 2-2.5 s (after 4-6 sessions).

#### 4.3.3 Phase 3: Learning to inhibit the Go response on Stop trials

In the third training step rats were introduced to the concept of Stop trials. Now, on 100% of trials, a Stop cue (5 kHz, ~75 dB tone) was presented simultaneously with the Go cue. This Stop cue indicated that rats should ignore the Go cue, abort the planned Go response, and instead stay in the nose poke port until a minimum stop wait time (WT) had passed and the Stop tone was turned off. The required WT was initially set to 250 ms and gradually increased with 50 ms per 25 correct trials, to a final value of 700 ms. Importantly, after successfully waiting in the nose poke port, rats were not allowed to press the lever before collecting a reward. Such erroneous lever presses, which were very rare (< 1% of trials), resulted in a 5 s timeout and omission of the reward. When rats could reliably inhibit the Go response and hold the nosepoke for a minimum of 700 ms (> 80% accuracy for ≥ 100 trials), they were moved to the final training phase (after 2-3 sessions).

#### 4.3.4 Phase 4: Stop Signal Task

In the final task rats were at first randomly presented with Go and Stop trials at a variable ratio. This ratio was 70:30 at the start of a session but varied over time based on the performance of a rat in both types of trials. All rats started with the following settings: maximum GoRT 0.8-1 s, maximum MT 2 s, minimum WT 0.4-0.5 s. Then, across several sessions, GoRT and WT were individually titrated in steps of 50 ms such that in the end both had the same value (between 0.4-0.7 s). MT on the other hand was gradually reduced to 1.2 s. We found that this MT value gives rats enough time to move to the lever after exiting the nosepoke port, but not so much time as to be able to e.g. check the reward port on their way to the lever. When the maximum GoRT and minimum WT were equal (after 11-14 sessions), the ratio between Go and Stop trials was fixed at 70:30, the maximum time to initiate a trial was reduced to 15 s, and session length was further reduced to 500 trials or 60 min, whichever came first. This prevented excessive variation in the number of trials rats completed during a session. After an additional 2-4 sessions, rats progressed to the full version of the SST that included SSDs. For each Stop trial, the SSD was randomly set to 0 ms or a value equal to mean GoRT - 250 ms, mean GoRT - 150 ms or mean GoRT - 50 ms. Here, mean GoRT was calculated before each session based on a rat’s performance during the previous five training sessions. On Stop trials where a rat initiated a Go response before presentation of the Stop cue (GoRT < SSD), that trial for the rat continued as if it was a Go trial, while still being recorded as a failed Stop trial for data analysis purposes e.g. (Beuk et al., 2014; Mayse et al., 2014; Schmidt et al., 2013). After introduction of SSDs, training continued until the whole cohort had reached stable baseline performance in the SST (15-23 sessions). During this period, for some rats, SSDs were occasionally omitted for 1-2 sessions. We found that this approach prevented rats from developing a strong bias towards either trial type.

#### 4.3.5 Behavioral pharmacology

The week before testing, rats were on 3 occasions habituated to an intraperitoneal (i.p.) saline injection. Pharmacological testing was performed according to a within-subject Latin square design, with at least 2 regular training sessions between test days. On each of the three test days, SB-334867 (NIDA, Research Triangle Park, NC) was dissolved in 2% dimethyl sulfoxide and then further diluted using a slightly heated solution of 10% 2-hydropropyl-β-cyclodextrin in sterile water. The resulting emulsion (or its vehicle) was immediately injected i.p. in a volume of 4 ml/kg, and the behavioral session started 30 min later.

### 4.4 Estimation of stop signal reaction time and calculation of other performance measures

All data were processed using custom-written MATLAB scripts. SSRTs were estimated with the recently-developed “modified integration method” (Mayse et al., 2014), a variation of the classical method for estimating the SSRT (Logan and Cowan, 1984; Logan, 1994). In short, first, *n* (where *n* equals the number of Stop trials in a session) random samples were drawn without replacement from the approximately 2.33*n* Go trials in a session. Next, the SSDs associated with the Stop trials in that session were subtracted from these *n* sampled GoRTs to create a new distribution of GoRTs re-aligned to would-be Stop signals. This sampling procedure was repeated 10,000 times to determine a conservative 99.9% (0.05–99.95%) confidence interval (CI) of the cumulative re-aligned GoRT distribution. Stop trial WTs were also aligned to the onset of the Stop signal and sorted to create a Stop trial cumulative WT distribution. Next, a WT cut-off was determined by finding the earliest timepoint at which the sorted Stop trial cumulative WT distribution dropped below the lower bound CI of the cumulative re-aligned GoRT distribution, with significant slowing of the Stop trial cumulative RT distribution present in at least ~0.15*n* consecutive Stop trials following that timepoint. The latter step guards against false positive discovery. The identified WT cut-off was then used to objectively separate Stop trials in which the rat failed to inhibit the Go response (Failed Stop errors) from Stop trials in which the rat after an initial successful inhibition of the Go response did not wait long enough (Waiting errors). Thus, the WT cut off determined the proportion of Failed Stop errors, *p*(Failed Stop), and Waiting errors, *p*(Waiting Error), among all Stop trials in a session. Finally, the mean of the re-aligned cumulative GoRT distributions was taken as the best estimate of the re-aligned cumulative GoRT distribution, and the estimated SSRT was defined as the timepoint in this distribution that corresponded to *p*(Failed Stop).

In addition to the above-mentioned three indices of Stop trial performance, we calculated one more measure of waiting impulsivity: percentage premature responding, i.e. trials during which the nosepoke port was exited during the hold period preceding Go cue presentation. We also calculated four measures of Go trial performance:: 1) *p*(Correct Go), the probability of correctly responding during Go trials (i.e. accuracy); 2) median GoRT, the median reaction time on correct Go trials; 3) ICV [standard deviation GoRT/mean GoRT] as a measure of Go response variability (e.g. (Bellgrove et al., 2004); and 4) median MT, the median time between exit of the nosepoke port and a lever press on correct Go trials. Task engagement and motivation were assessed by calculating the number of trials/min executed, percentage of trials omitted (trials that were not initiated by the rat within 15 s of collecting the reward on the previous trial), and median feeder latency – the median time to collect a reward following correct completion of a Go or Stop trial (e.g. (Bari and Robbins, 2013; Humby et al., 2013). Finally, measures of trial type discrimination and overall response bias derived from signal detection theory measures were calculated (e.g. (Macmillan and Creelman, 2005; Stanislaw and Todorov, 1999): Sensitivity index (d’) = *Z*(*p*[Correct Go]) – *Z*(*p*[Failed Stop] + *p*[Waiting Error]) and response bias (c) = 0.5 ^*^ (*Z*(*p*[Correct Go]) + *Z*(*p*[Failed Stop] + *p*[Waiting Error])), where *Z*(*p*), *p* ∈ [0,1], is the inverse of the cumulative Gaussian distribution.

### 4.5 Statistical analysis

An average baseline performance of each rat was calculated by averaging data from the 3 training sessions that immediately preceded the three test days. Rats were excluded from further analysis if their baseline performance violated one or more of the following criteria: 1) average *p*(Correct Go) > 0.7 with at most 1 day *p*(Correct Go) < 0.7; 2) SSRT > 50 ms; 3) *p*(Failed Stop) increasing across increasing SSDs; 4) *p*(Failed Stop) and *p*(Waiting Error) < 0.3 when SSD = 0 ms. In total, six rats were excluded this way; two for violating criterion 1, two for violating criterion 4, and a final two for violating criteria 1 and 4.

Statistical analysis for all data was performed using NCSS 11 (NCSS, LLC., Kaysville, UT). Data were first checked for normal distribution and homogeneity of variance. Normal distribution of the data was verified by Shapiro-Wilk testing and visual inspection. Data for three parameters (premature response rate, p(Waiting Error), omission rate and trial execution rate) were found to fail normality testing, and were therefore square root-transformed before further analysis. In the result section untransformed data is displayed for all parameters, for ease of interpretation. Data were then subjected to repeated measures analyses of variance with drug treatment and SSD (for p(Failed Stop) and p(Waiting Error) only) as within-subject variables. Homogeneity of variance across groups was determined using Mauchly’s tests for equal variances. In case of violation of homogeneity, Geisser-Greenhouse epsilon (ε) adjusted degrees of freedom were applied and the resulting more conservative F-values and probability values depicted (unadjusted degrees of freedom are depicted in the results section for easier interpretation). In case of statistically significant main effects (*p* < 0.005 was used as cut-off to correct for the large number of behavioral parameters assessed), further *post hoc* comparisons were conducted using Bonferroni multiple comparison tests with *p* < 0.05 indicating statistically significant effects. All graphs were produced using GraphPad Prism version 6 (GraphPad Software, San Diego, CA).

## Author contributions

JW, MHJ and GAJ were responsible for the overall study design. JW and MHJ performed the experiments. JW analyzed the data. JW and MHJ wrote the initial versions of the manuscript, while JW, MHJ and GAJ wrote the final version of the manuscript.

## Acknowledgements

The authors would like to thank Shayna L. O’Connor and Sasha (A.L.) Taylor for their help with the performing the experiments, and Dr. Shih-Chieh Lin for help with implementing the new methods for stop signal reaction time estimation. We also thank Ms. Nupur Jain for her general assistance.

## Abbreviations

5-CSRTT: five choice serial reaction time task
DMH/PF: dorsomedial hypothalamus and perifornical area
GoRT: Go trial reaction time
ICV: intra-individual coefficient of variation
LH: lateral hypothalamus
MT: movement time
OxR1 and OxR2: orexin-1 and orexin-2 receptors
SB: SB-334867, orexin-1 receptor antagonist
SSD: stop signal delay
SSRT: stop signal reaction time
SST: stop signal task
WT: stop trial wait time

